# Community origins and regional differences highlight risk of plasmid-mediated fluoroquinolone resistant Enterobacteriaceae infections in children

**DOI:** 10.1101/301457

**Authors:** Latania K. Logan, Rachel L. Medernach, Jared R. Rispens, Steven H. Marshall, Andrea M. Hujer, T. Nicholas Domitrovic, Susan D. Rudin, Xiaotian Zheng, Nadia K. Qureshi, Sreenivas Konda, Mary K. Hayden, Robert A. Weinstein, Robert A. Bonomo

**Author notes:** **Corresponding Author:** Latania K. Logan, Rush University Medical Center 1620 W. Harrison, Suite 951 Jelke, Chicago, IL 60612. **Financial Disclosure:** The authors have no financial disclosures relevant to this article.

## Abstract

**Background:** Fluoroquinolones (FQs) are uncommonly prescribed in children, yet pediatric multidrug-resistant (MDR)-Enterobacteriaceae (Ent) infections often reveal FQ resistance (FQR). We sought to define the molecular epidemiology of FQR and MDR-Ent in children.

**Methods:** A case-control analysis of children with MDR-Ent infections at 3 Chicago hospitals was performed. Cases were children with third-generation-cephalosporin-resistant (3GCR) and/or carbapenem-resistant (CR)-Ent infections. PCR and DNA analysis assessed *bla* and plasmid-mediated FQR (PMFQR) genes. Controls were children with 3GC and carbapenem susceptible-Ent infections matched by age, source and hospital. We assessed clinical-epidemiologic predictors of PMFQR Ent infection.

**Results:** Of 169 3GCR and/or CR Ent isolates from children (median age 4.8 years), 85 were FQR; 56 (66%) contained PMFQR genes. The predominant organism was *E. coli* and most common *bla* gene *bla* _CTX-M-1 group_. In FQR isolates, PMFQR gene mutations included *aac6’1b-cr, oqxA/B, qepA*, and *qnrA/B/D/S* in 83%, 15%, 13% and 11% of isolates, respectively. FQR *E. coli* was often associated with phylogroup B2, ST43/ST131. On multivariable analysis, PMFQR *Ent* infections occurred mostly in outpatients (OR 33.1) of non-black-white-Hispanic race (OR 6.5). Residents of Southwest Chicago were >5 times more likely to have PMFQR-*Ent* infections than those in the reference region, while residence in Central Chicago was associated with a 97% decreased risk. Other demographic, comorbidity, invasive-device, antibiotic use, or healthcare differences were not found.

**Conclusions:** The strong association of infection with MDROs showing FQR with patient residence rather than with traditional risk factors suggests that the community environment is a major contributor to spread of these pathogens in children.

## Introduction

Multi-drug resistant (MDR) Enterobacteriaceae infections are associated with significant morbidity and mortality and are an emerging problem in children in the US during the last decade (1, 2). Globally, this has been attributed mainly to the rise in extended-spectrum beta lactamase (ESBL) producing Enterobacteriaceae (ESBL Ent) and carbapenem-resistant Enterobacteriaceae (CRE) (3, 4). Increasing resistance to several other classes of antibiotics was found in these studies; including to the fluoroquinolones, a class of antibiotics with limited indications for use in children (1, 2).

The CTX-M-ESBL harboring *Escherichia coli* are among the most common multi-drug resistant organisms (MDROs), often possessing resistance genes to other important antibiotic classes including aminoglycosides, fluoroquinolones, tetracyclines, and trimethoprim-sulfamethoxazole (5). In adults, fluoroquinolone resistance in (FQR) Gram-negative bacteria has been linked to chromosomal and to plasmid-mediated resistance mechanisms and is thought to be associated mostly with the dramatic increase in use of these antibiotics during the 1980s(6).

The reasons for increased FQR in children are unclear, and studies assessing the resistance determinants associated with the FQR phenotype in Enterobacteriaceae recovered from children are limited(7); therefore, we examined this population to determine which children had a higher likelihood of infections with organisms resistant to both beta-lactam and fluoroquinolone antibiotics.

We determined the genetic basis of FQR in beta-lactamase producing Enterobacteriaceae isolates from children cared for by multiple centers in the Chicago area and analyzed a subgroup of children with infections with similar resistance determinants, namely those due to isolates containing genes encoding plasmid-mediated fluoroquinolone resistance (PMFQR) and ESBL mediated resistance to determine genotypes, host factors, and exposures leading to infection with MDR Enterobacteriaceae strains. We hypothesized that because of differences in healthcare delivery in urban settings, acquisition of PMFQR in children would be linked to geographic location and have environmental influences and community origins. Our analysis of host factors and exposures revealed that there are genetic and geospatial links to MDR in this pediatric population.

## METHODS

### Study Setting

Hospital A contains a 115-bed children’s hospital within a tertiary care academic medical center which has a mother-newborn infant unit, pediatric and psychiatric wards, and cardiac, pediatric and neonatal intensive-care units (PICU and NICU). Hospital B has 288 beds and is a free-standing children’s academic medical center that provides complex quaternary services, such as pediatric organ and bone marrow transplantation. Hospital C is a 125-bed children’s hospital within an academic medical center, and contains general pediatrics and newborn infant wards, as well as a PICU and NICU. All of the participating centers are within metropolitan Chicago.

### Descriptive Study Design

### Study Population

This study included patients aged 0 to 18.99 years who had clinical cultures positive for Enterobacteriaceae with 3GCR or CR and the suspected presence of a beta-lactamase gene based on clinical laboratory testing. Additionally, isolates found to be concomitantly resistant to FQR were further characterized. Infections were diagnosed between January 1, 2011 and December 31, 2014 and only the first infection per patient was included. The study was approved by the institutional review boards of the three participating institutions and need for informed consent was waived.

### Testing of Antibiotic Susceptibility in Enterobacteriaceae

The Hospitals A-C microbiology laboratories phenotypically analyzed presumed ESBL Ent, AmpC Ent and CR isolates via the Vitek 2 microbial identification system (*bioMérieux, Athens, GA)* or by the MicroScan WalkAway system (Siemens Healthcare Diagnostics, Tarrytown, NY). Screening for ESBL production involved testing with one or more of the following agents: aztreonam, ceftazidime, ceftriaxone, cefotaxime or cefpodoxime, based on guidelines of the Clinical and Laboratory Standards Institute (CLSI) (8). ESBL production was confirmed on the automated instruments or by disk diffusion assays (BBL; Becton, Dickinson and Company, Sparks, MD) or by measuring minimum inhibitory concentrations (MICs) of ceftazidime and cefotaxime in the presence and absence of clavulanic acid. A measurement of an increase in disk zone diameter of > 5 mm or a 4-fold reduction in the MIC of ceftazidime or cefotaxime in the presence of clavulanic acid served as confirmation of the ESBL phenotype (8).

The carbapenemase phenotype, per Centers for Disease Control and Prevention (CDC) criteria, included isolates that were non-susceptible to all 3GCs (cefotaxime, ceftazidime, or ceftriaxone) and resistant to one or more carbapenem (imipenem, meropenem, doripenem, or ertapenem) (9). Carbapenemase production was phenotypically confirmed by MBL E-test (bioMérieux, Athens, GA) or Modified Hodge Test, as appropriate (10).

### Determination of Beta-Lactam Resistance Mechanisms

Genomic DNA was extracted and purified from isolates using the DNeasy Blood & Tissue Kit (QIAGEN, Inc., Valencia, CA). To evaluate for the presence of *bla* genes in isolates, a DNA microarray based assay was performed (Check-Points, Check-MDR CT101 kit; Wageningen, The Netherlands). The CT101 microarray based assay can detect the following *bla* groups: CTX-M-1 group, CTX-M-2 group, CTX-M-8 and -25 group, CTX-M-9 group, SHV WT and SHV-type ESBL, TEM wild-type, and TEM-type ESBL, plasmid based AmpC cephalosporinases (pAmpC) (CMY II, ACC, FOX, DHA, ACT/MIR) and carbapenemases (KPC and NDM) (11). When isolates were *bla* negative by the CT101 assay, a broader DNA microarray, (Check-Points, Check-MDR CT103XL kit) was performed. The CT103XL assay can detect the presence of additional ESBL genes (VEB, PER, BEL, GES) and carbapenemase genes (GES, GIM, IMP, SPM, VIM, and OXA-23, -24/40, -48, and -58) (12). The assays were performed as described in our laboratory previously (13).

### Analysis of Determinants Yielding Fluoroquinolone Resistance

To investigate the presence of FQR determinants in MDR Enterobacteriaceae isolates, we analyzed the quinolone resistance-determining region (QRDR) located on the bacterial chromosome and assessed for PMFQR in strains found FQR by CLSI standards (8). Briefly, genus specific assays for mutations in *gyrA* and *parC* genes of the QRDR (in *E. coli, Klebsiella sp.*, and *Enterobacter sp.*) and for PMFQR were performed by polymerase chain reaction (PCR) and deoxyribonucleic acid (DNA) sequencing of amplicons (6). Extraction of genomic DNA followed by amplification and sequencing were performed using primers and methods as previously described (14, 15). Specific PMFQR genes screened include *qnrA, qnrB, qnrD, qnrS, qepA, oqxA* and *oqxB* and *aac6’-Ib-cr* and represent transmissible elements reported in Enterobacteriaceae (16).

### Multilocus Sequence Typing (MLST)

Per protocol, eight *E. coli* housekeeping genes (*dinB, icdA, pabB, polB, putP, trpA, trpB and uidA*) and seven *Klebsiella* species (sp.) housekeeping genes *(rpoB*, *gapA, mdh, pgi, phoE, tonB, infB)* were amplified and sequenced as in prior studies (13). Alleles and sequence types (ST) were assigned for select isolates of varying genotypic profiles by the Pasteur MLST scheme (http://www.pasteur.fr/recherche/genopole/PF8/mlst/).

### Analysis of Plasmid Replicon Types and Phylogenetic Grouping

*E.coli* were assigned to four major phylogenetic groups (A, B1, B2 and D) using a well-established multiplex PCR-based method (17). Plasmids were typed, in select isolates with varying genotypic profiles, based on incompatibility groups corresponding to the nomenclature assigned by Carattoli *et al*. (18).

### Analytic Study Design

We used a retrospective case–control study design to assess factors associated with infection due to beta-lactam, FQ resistant isolates in which we had detected a PMFQR gene. We chose to analyze FQR in detail because this class of antibiotics is uncommonly used in children, yet 50% of the isolates between 2011 and 2014 were FQR.

We selected as controls, children with infections due to bacteria susceptible to the antibiotics of interest. Specifically, the control group included children with infections that were susceptible to 3GC, carbapenem and FQ antibiotics to understand differences between children who acquired plasmid-mediated MDR Enterobacteriaceae infections and those who did not.

Only patients with clinical infections were included, as determined by study investigator case review and/or using standard criteria defined by the CDC National Healthcare Safety Network (19). Children serving as control subjects were identified using hospital electronic laboratory records (ELRs). Control patients were matched approximately 3:1 to the cases by age range, hospital, and specimen source.

### Covariates

Several variables were analyzed as potential factors associated with FQR Ent infection based on known associations for acquisition in adults including (1) demographics (age, gender, race/ethnicity); (2) comorbid conditions (as defined by ICD-9 codes); (3) recent inpatient and outpatient healthcare exposures, including hospitalization and/or procedures in the previous 30 days; (4) all recent antibiotic exposures in the 40 days prior to culture (32); (5) presence, number, and type of invasive medical devices; and (6) the impact of location of patient residence in the Chicago area as assessed by dividing the metropolitan area into 7 regions using zip code level data, which included Chicago proper and its suburban areas (i.e. Northwest side and Northwest Suburbs, Southwest side and Southwest Suburbs, etc.). An eighth region included patients from other parts of Illinois or from other states.

### Statistical Analysis

Case and control groups were examined for differences using parametric or non-parametric tests as appropriate for categorical and continuous variables; P≤0.05 was considered statistically significant unless otherwise specified. Variables with p<0.1 on bivariate analysis were included in multivariable analysis. Stepwise multiple logistic regression was used to assess the multivariable relationship between the covariates and the groups. The final multivariable logistic regression model included the simplest model with significant covariates (p<0.05) from the stepwise selection process, with PMFQR Ent infection as the outcome variable. The simplest model was chosen based on a relatively small sample size and the effect of variables in the model. All analyses were performed in SAS 9.4 (SAS Institute, Cary, NC, USA).

## RESULTS

### Composition of Fluoroquinolone Resistance (FQR) Genes in Enterobacteriaceae

We assessed 169 *bla*-producing Ent isolates between 2011 – 2014 from Hospitals A, B, and C for the presence of FQR (Table 1). Of 169 Ent isolates, 85 (50%) were FQR of which 82 (96.4%) were available for further testing. The median age of children with FQR-Ent infections was 4.8 years. The predominant organism was *E. coli*, 65/82 (79%), and the predominant *bla* genotype found associated with FQR in Ent was *bla*_CTX-M-1-group_ in 62% of cases. Within *E. coli*, FQR was most often associated with phylogroup B2 and ST43 (Pasteur scheme)/ST131 (Achtman scheme) harboring *bla*_CTX-M-1-group_ in 47/63 (75%) cases.

**TABLE 1.**
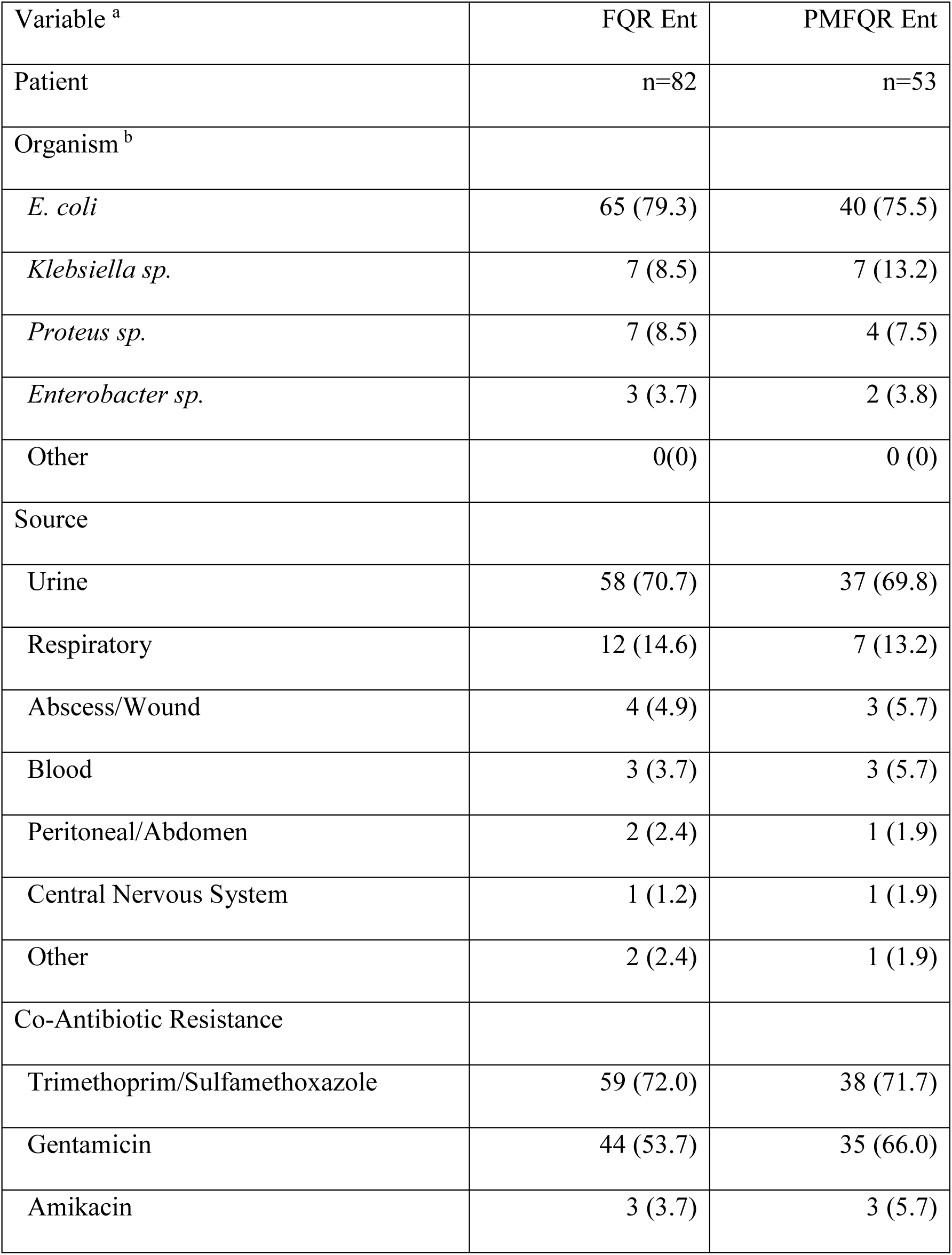
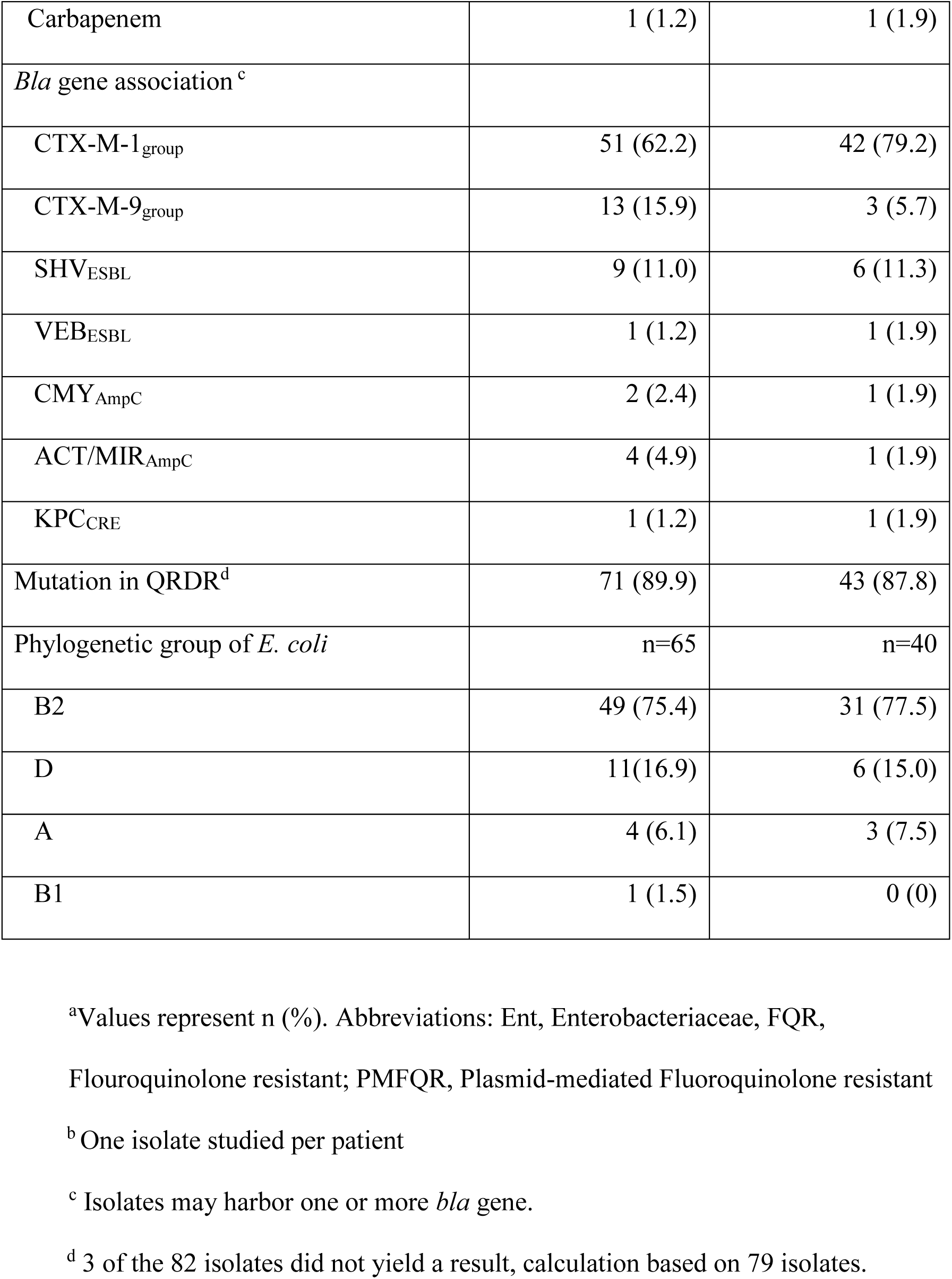
CHARACTERISTICS OF FQR AND PMFQR ENTEROBACTERIACEA

FQR isolates were further characterized to understand resistance determinants associated with FQR in pediatric Ent isolates. Chromosomal mutations of the QRDR (*gyrA*/*parC*) were present in 71/79 (89.9%) of FQR isolates by DNA sequence analysis. Three isolates did not yield results. PMFQR genes were detected by PCR in 56/82 (66%); 53 (95%) were available for further analysis. The median case patient age was 6 years. PMFQR genes included *aac 6’1b-cr, oqx A/B, qepA*, and *qnr A/B/D/S* in 83%, 15%, 13% and 11% of isolates, respectively. PMFQR was found in combination with *gyrA* and/or *parC* mutations in 43/49 (88%) isolates, which is associated with high level resistance. The predominant *bla* genotype found associated with PMFQR was *bla* _CTX-M-1-group_ in 76%, followed by *bla*SHV ESBL associations in 11%. Almost all (98%) PMFQR Ent were multi-drug resistant, e.g. resistant to ≥3 antibiotic classes.

### Analysis of Factors Associated with PMFQR Enterobacteriaceae Infections in Children

The 53 cases of PMFQR Ent infection were matched by age range, hospital, and culture source to 131 controls with antibiotic-sensitive Ent infections. Significant factors associated with PMFQR Ent infection on bivariate analysis included: *E. coli* infection, race/ethnicity, infection diagnosed in an outpatient clinic, history of quinolone use, and residence in the Southwest region (hereafter referred to as the high-risk region) comprised of southwest Chicago and the southwestern Chicago suburbs (Table 2). Children with PMFQR infection were less likely to have infection with *Enterobacter* sp., a central venous catheter, neonatal intensive care unit admission at the time of infection diagnosis, or residence outside the “high-risk” region (comprised of the downtown Chicago area, near North side, Chicago loop, and North Chicago). Case-control differences in comorbid conditions; presence of respiratory, gastrointestinal, genitourinary, or overall count of foreign bodies; or recent prior healthcare exposure were not found.

**TABLE 2.** BIVARIATE ANALYSIS OF DEMOGRAPHICS AND FACTORS ASSOCIATED WITH PMFQR ENTEROBACTERIACEAE INFECTION

| Characteristic<sup>a</sup> | PMFQR<sup>b</sup> Infection | Non-PMFQR<br>Infection<sup>c</sup> | p value |
| --- | --- | --- | --- |
| Patient | n=53 | n=131 |  |
| Location at Diagnosis |  |  | <0.0001 |
| Inpatient, non ICU | 21 (39.6) | 46 (35.1) |  |
| Outpatient Clinic | 16 (30.2) | 7 (5.3) |  |
| Emergency Room | 4 (7.6) | 23 (17.6) |  |
| Pediatric ICU | 11 (20.8) | 37 (28.2) |  |
| Neonatal ICU | 1 (1.9) | 18 (13.7) |  |
| Region of Residence<sup>d</sup> |  |  | 0.047 |
| Downtown | 1 (1.9) | 19 (14.5) |  |
| Northwest | 6 (11.3) | 14 (10.7) |  |
| Far North | 14 (26.4) | 25 (19.1) |  |
| West | 12 (26.4) | 36 (27.5) |  |
| Southwest | 10 (18.9) | 8 (6.1) |  |
| South | 2 (3.8) | 12 (9.2) |  |
| Far South | 3 (5.7) | 8 (6.1) |  |
| Other IL/Other states | 3 (5.7) | 9 (6.9) |  |
| Recent Health Care |  |  | 0.406 |
| Inpatient Care | 12 (22.6) | 31 (23.7) |  |
| Outpatient Care<sup>e</sup> | 31 (58.5) | 64 (48.9) |  |
| No Recent Care | 10 (18.9) | 36 (27.5) |  |
| Cephalosporins<sup>f</sup> | 15 (28.9) | 33 (25.4) | 0.63 |
| Fluoroquinolones<sup>g</sup> | 6 (11.5) | 3 (2.3) | 0.01 |
| TMP/SMX<sup>h</sup> | 8 (15.4) | 18 (13.9) | 0.79 |
| Central venous line | 9 (17.0) | 44 (33.6) | 0.025 |
| Gastrointestinal | 13 (24.5) | 45 (34.4) | 0.195 |
| Genitourinary | 14 (26.4) | 37 (28.2) | 0.802 |
| Respiratory | 11 (20.8) | 38 (29.0) | 0.253 |

<sup>a</sup> Values represent n (%) unless otherwise indicated.
<sup>b</sup> Abbreviation PMFQR, Plasmid-mediated Fluoroquinolone Resistance
<sup>c</sup> Non-PMFQR Infection were children with infections due to Enterobacteriaceae sensitive to extended spectrum cephalosporin and fluoroquinolone antibiotics.
<sup>d</sup> Region of residence includes city region and neighboring suburbs, except for “Downtown”, which includes downtown, near North side, loop, and Northside, all within city limits. “Other” includes other cities in Illinois not neighboring Chicago, and patients from other states.
<sup>e</sup> Outpatient care includes care outside of routine child care visits and outpatient procedures.
<sup>f</sup> Includes previous exposure to extended-spectrum cephalosporins (ceftriaxone, ceftazidime, cefotaxime)
<sup>g</sup> Includes previous exposure to ciprofloxacin and levofloxacin
<sup>h</sup> Includes previous exposure to TMP-SMX, Trimethoprim-Sulfamethoxazole

We did not find evidence of significant effect modification during the model building stages nor did we find evidence of significant confounding; therefore, no additional covariates were added back to the final model after the stepwise selection process was completed, and the simplest model was used in the final regression model.

On multivariable analysis (Table 3), having infection diagnosed in the outpatient clinic setting was significantly associated with PMFQR Enterobacteriaceae infection (OR=33.1; 95% CI 7.1, 162.8; p<0.001). Being of a race or ethnicity other than white, black, or Hispanic was significantly associated with PMFQR Enterobacteriaceae infection (OR 6.5; 95% CI 1.7, 24.3; p=0.006). Interestingly, among children with Enterobacteriaceae infections, those residing in southwestern region of Chicago had more than five times the odds of having a PMFQR infection compared to those living in the reference West Chicago region (OR 5.6; 95% CI 1.6, 19.2; p=0.006) after controlling for race and healthcare setting. In contrast, for children who resided in the downtown region, there was a 97% decrease in the odds of PMFQR infection in those living in this region compared to those residing in reference West Chicago region (OR 0.03; 95% CI .002, 0.33; p=0.005). No other regional associations were found.

**TABLE 3.** MULTIVARIABLE ANALYSIS OF FACTORS ASSOCIATED WITH PMFQR ENTEROBACTERIACEAE INFECTIONS IN CHILDREN

| Associated Factor with PMFQR infection<sup>a</sup> | OR | 95% CI | p value |
| --- | --- | --- | --- |
| Outpatient Clinic Location at time of infection<br>diagnosis | 33.9 | 7.08, 162.8 | <0.001 |
| Race not white, black or Hispanic | 6.5 | 1.70, 24.3 | 0.006 |
| Home Residence in Southwest Region (SW Chicago<br>and SW Suburbs) | 5.6 | 1.6, 19.2 | 0.006 |
| Home Residence in Downtown Region (Near North<br>Side, Loop, Northside) | 0.03 | 0.002, 0.33 | 0.005 |

<sup>a</sup> Abbreviations, SW, Southwest; Loop, Chicago Loop.
<sup>b</sup> Reference Region, West Region.

To ensure these regional associations were unique to PMFQR containing isolates, we ran two additional case-control analyses specifically assessing whether regional differences within the Chicago metropolitan area were seen with 1) ESBL-producing strains that were sensitive to fluoroquinolones; and 2) *bla*_CTX-M-1-group_ ESBL-producing isolates (related to circulation of ST131 clonal *E. coli* strains). Regional differences in acquisition were not found in either analysis (data not shown).

## DISCUSSION

Multi-drug resistant Enterobacteriaceae are a growing concern globally. Much of the propagation and spread of these organisms has been related to the ST131 *E. coli* strains and to high-risk clones containing IncFII plasmids and other genetic structures, such as transposons, integrons, and insertion sequences associated with multiple antibiotic resistance gene cassettes (20). Occasionally, beta-lactamase and other antibiotic resistance genes are transferred horizontally (7).

In our pediatric patients there was a predominance of ST131 *E. coli* harboring *bla* _CTX-M_. We hypothesize that there is significant horizontal gene transfer between genera. This is very worrisome from a public health perspective since children, once colonized with MDR Enterobacteriaceae can remain colonized for months to years and could serve as reservoirs and “silent disseminators” of MDROs (21). Interestingly, the community focus of the MDROs in children is in stark contrast to the epidemiology of these bacteria in adults in Chicago, where MDR Enterobacteriaceae acquisition is highly linked to residence in long-term care facilities and to interfacility transfer (22).

We found striking residential differences for children infected with PMFQR containing Enterobacteriaceae (PMFQR Ent) compared to children infected with antibiotic sensitive strains. PMFQR Ent contain plasmid-based resistance genes to fluoroquinolones, an antibiotic uncommonly used in children. In one Chicago region, the Southwest region, there was a substantial increase in odds of PMFQR Ent infection, and in the Downtown region, there was a significant decrease in the likelihood of PMFQR Ent infection. Our study included 3 major pediatric centers, none of which (and no major medical centers for children) is located in the “high-risk” Southwest region, yet all three centers diagnosed and treated patients with PMFQR Ent infections from that region. In contrast, Hospital B, the largest hospital in the region specifically dedicated to the care of children, is located in the downtown region, and therefore services many children residing in that area; yet this was the area of “lowest risk” for PMFQR Ent infection. This may reflect linkage to *bla* _CTX-M_ harboring plasmids which are endemic in some communities, but the reservoirs are currently undefined.

Interestingly, we found that children with PMFQR Ent infection were more likely to present in the outpatient clinic setting than were those with antibiotic sensitive Ent infections – the opposite of expectation that MDROs are healthcare linked and suggesting that PMFQR Ent infections were community acquired and less severe.

A strong additional association on multivariable analysis was the higher likelihood of PMFQR Ent infection in those of non-white, non-black, and non-Hispanic race. This risk was statistically independent of residence, and strikingly none of the children located in the “high risk” southwest region with PMFQR Ent infection were of race “other”, supporting the strong independence of these 2 risk factors. Due to the retrospective nature of the study, we were unable to gather further data on “other” race or ethnicity. We did not have travel data for the majority of children, although it is well documented that travel to certain countries can be associated with high rates of ESBL Ent acquisition, particularly in South and Southeast Asia (23, 24). It is also well described that there is an increased risk of colonization of household members after return of the traveler who first acquired an ESBL Ent.(25)

We did not find significant differences in comorbidities between cases and controls. This puts further importance on evaluating environmental sources for plasmid-possessing antibiotic resistance genes (26). Community-based environmental influences would include higher exposure risks in certain communities due to certain foods, livestock, animals, water sources, fertilizer, soil, and vegetation (27). For example, if there is a link to food exposures, such as restaurant chains that cook with high saturated fats, and additionally serve food animals that are fed antibiotics and hormones for growth effects, this exposure would increase the risk of acquisition of antibiotic-resistant bacteria, as well as obesity (28). This in turn increases the risk of other diseases such as cardiovascular disease and diabetes (29). Some of the PMFQR genes, for example, oqxA and oqxB are multidrug efflux pumps named for their resistance to olaquinadox, which is used as a growth promoter on pig farms (30). We did not have data on companion pets for the majority of children and were unable to examine this association, though this may play an important role in acquisition of resistant pathogens (9).

Studies in our region and nationally have suggested that an increased risk of exposure to antibiotics in children (31), as well as to antibiotic resistant bacteria, may be related to socioeconomic status and race (32). While we did assess race, we did not formally compare differences between socioeconomic factors in the regions, as we did not have street or neighborhood level data on infected patients. However, in a general comparison of regional zip codes using Illinois census data, we did not find overall differences in the socioeconomic status of the “high risk” southwest region and neighboring regions such as the south and west regions. We did not have data on antibiotic usage in adults within the household, which could also increase acquisition of resistant pathogens.

We recognize that our study has limitations. This was a retrospective study designed to determine mechanisms of antibiotic resistance in Enterobacteriaceae recovered from children cared for at three centers in a single metropolitan area; this may potentially impact generalizability to other regions. Additionally, a plasmid-based origin of the recovered antibiotic-resistance genes is suggested by our DNA sequence analysis results, yet it is possible that some of these genes represent chromosomal resistance mechanisms. However, subsequent plasmid-replicon typing and DNA sequence analysis for a subgroup of bacteria support our findings of the DNA microarray. Our sample size was relatively small, which may allow for selection bias; however, the pooling of multicentered data from institutions of differing types serving diverse populations throughout the 3rd largest metropolitan area in the U.S. potentially lessens this bias. The smaller sample sizes in pediatric studies are related to the overall low prevalence of these organisms in children in most U.S. areas (1-3%), including in the Chicago and the Midwest region (21, 23, 26), although national trends indicate an increase in prevalence of these menacing organisms in pediatric populations during the last decade, suggesting they are an emerging threat that needs further evaluation (7).

In conclusion, we found that there is significant complexity and diversity in the determinants associated with beta-lactam and fluoroquinolone antibiotic resistance in children, and that pediatric MDR Enterobacteriaceae exhibited differences when compared to descriptions of strains circulating in adult patients in a region where such infections are endemic. We also describe, for the first time, the impact of residence on infection with MDR Enterobacteriaceae in children located in the same geographic area, however the reservoirs remain undefined. Future studies should focus on further molecular characterization of circulating strains and the environmental influences associated with these differences in regional acquisition. We anticipate that an imminent threat of the “silent dissemination” of multi-drug resistant Enterobacteriaceae in community settings is occurring in children. Local, federal, and international programs dedicated to this serious problem must focus on halting the spread of these menacing pathogens in our most vulnerable population, children.

## Acknowledgements

We gratefully acknowledge the contribution of the late Dr. Paul Schreckenberger to this work. We thank the microbiology laboratories of the participating institutions for providing isolates for this study. We thank Kendrick Reme and Lynika Strozier of the Logan Laboratory and Pamela Hagen, Jane Stevens, Joyce Houlihan, Kathleen McKinley, Violeta Rekasiu, Cindy Bethel, and Donna Carter of participating institutions for collection, shipping, and cultivation of organisms. We thank the team of curators of the Institut Pasteur MLST and whole-genome MLST databases for curating the data and making them publicly available at http://bigsdb.web.pasteur.fr/. We report no conflicts of interest relevant to this study.

The content is solely the responsibility of the authors and does not necessarily represent the official views of the National Institutes of Health or the Department of Veterans Affairs.

## Funding

This work, including the efforts of Latania K. Logan, was funded by the National Institute of Allergy and Infectious Diseases, National Institutes of Health (NIH) (K08AI112506). This work, including the efforts of Robert A. Bonomo, was funded by the National Institute of Allergy and Infectious Diseases, National Institutes of Health (NIH) (R01AI072219, R01AI063517, and R01AI100560). R.A.B. is also supported by the Department of Veterans Affairs Research and Development under award number I01BX001974, VISN 10 Geriatrics Research, Education and Clinical Center.

## Financial Disclosure

The authors have no financial disclosures relevant to this article.

## Conflicts of Interest

The authors have no conflicts of interest relevant to this article.

## Notes

**Funding Source:** This work was directly supported by National Institutes of Health - National Institute of Allergy and Infectious Diseases grant K08AI112506 (Dr. Logan) and grants R01AI072219, R01AI063517, and R01AI100560 (Dr. Bonomo). This work was also supported by the Department of Veterans Affairs Research and Development under award number I01BX001974, VISN 10 Geriatrics Research, Education and Clinical Center (Dr. Bonomo).

